# Episodic, Semantic, Pavlovian, and Procedural Cognitive Maps

**DOI:** 10.1101/161141

**Authors:** José J. F. Ribas Fernandes, Clay B. Holroyd

## Abstract

Current theories of planning associate the hippocampus with a cognitive map, a theoretical construct used to predict the consequences of actions. This formulation is problematic for two reasons: First, cognitive maps are traditionally conceptualized to generalize over individual episodes, which conflicts with evidence associating the hippocampus with episodic memory, and second, it fails to explain seemingly non-hippocampal forms of planning. Here we propose a novel theoretical framework that resolves these issues: each long-term memory system is a cognitive map, predicting consequences of actions based on its unique computational properties. It follows that hippocampal maps are episode-based and that semantic, procedural, and Pavlovian memories each implement a specialized map. We present evidence for each type of map from neuropsychology, neuroimaging and animal electrophysiology studies.

## Main Text

In recent years, the cognitive neuroscience of **planning** (see Glossary) has been the subject of intense research interest. As defined within formal theories of **model-based reinforcement learning** (Box 1) (Daw & Dayan, 2014; Daw, Niv, & Dayan, 2005; Dolan & Dayan, 2013), planning is a process in which a **model** of a decision problem is used to calculate the value of actions before the actions are taken, and then deciding which actions to execute based on the results (Geffner & Bonet, 2013). The model represents the set of **transitions** and rewards, providing a formal means for predicting the consequences of actions. For example, a model could provide an answer to the questions, “Where do I end up when I take bus 15 from campus?” (a transition) and “How cheap and fast is bus 15?” (a reward). Crucially, models are usually conceived to be **abstract** or instance-less: they answer the general question, “What happens when I take bus 15?”, rather than the specific question, “What happened when I took bus 15 last Thursday at 5 p.m.?” The process of using a model to compare decision alternatives is referred to as **search**, a concept that aligns with the ideas of simulation, forethought, future-thinking, search, prospection, vicarious trial-and-error and **preplay** in other literatures (Michaelian, Klein, & Szpunar, 2016; Todd, Hills, & Robbins, 2012).

Decades prior to the introduction of model-based reinforcement learning, Edward Tolman proposed an analogous idea called a **cognitive map** based on his own empirical work (Tolman, 1948). Like models, cognitive maps refer to internal representations of latent causal relationships between actions and states that allow for search. Subsequent research has firmly linked cognitive maps and models with the **hippocampus** (Box 2) (Wikenheiser & Redish, 2015). However, this putative relationship raises a challenging problem: The function of the hippocampus as a cognitive map is incompatible with its more established role in **episodic** processing (Fig.1A) (Box 2) (Bendor & Spiers, 2016). In particular, episodic memories encode individual events associated with specific times and places, rather than abstract representations of these events.

**Figure 1.**
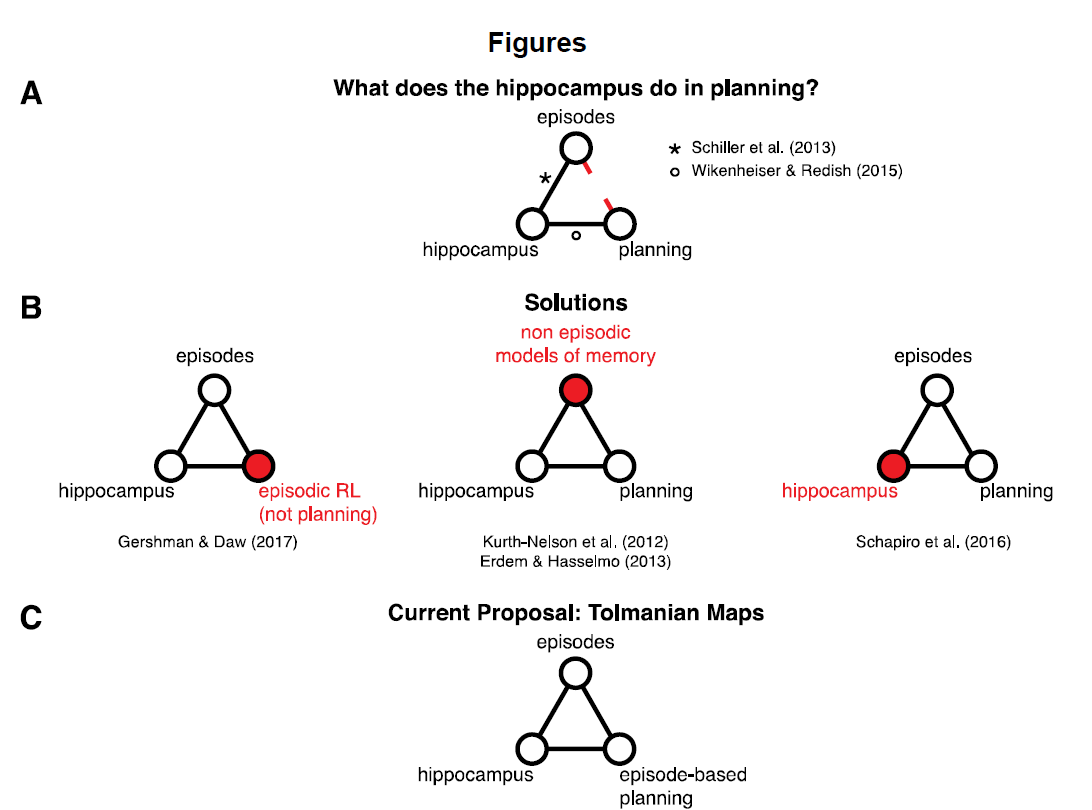
Hippocampus: Planning or Episodic Memory? *A: What does the hippocampus do in planning?* The idea that that the hippocampus implements a cognitive map that abstracts across instances, i.e., is “instance-less”, conflicts with its more established role in the formation and retrieval of episodic memories (2016). This discrepancy is reflected in diverging perspectives, rarely acknowledged, in review articles about hippocampal function that describe either the relationship between the hippocampus and episodes (Schiller et al., 2015), or the relationship between the hippocampus and planning (Wikenheiser & Redish, 2015), but not both (broken red connection). *B: Solutions.* Existing solutions change one element of this triad between the hippocampus, planning and episodic memory (red filled circles) in order to accommodate the data of the other 2 elements (open circles). The hippocampus has been argued to re-evaluate past episodes, which appears like planning but is not (Gershman & Daw, 2017) (left panel). Models of planning have been proposed to rely on properties of episodic memory and hippocampal dynamics that are not actually instance-based (Erdem & Hasselmo, 2012; Hopfield, 2010; Kurth-Nelson et al., 2012) (middle panel). Separate episodic and statistical functions – the latter of which underlie an abstract cognitive map – have been attributed to segregated hippocampal subregions (Schapiro, Turk-Browne, Botvinick, & Norman, 2016), leaving open the question of how the two processes can be reconciled in a single function (right panel). Therefore, the interrelationship between the hippocampus, planning, and episodic memories remains undetermined. *C: Current proposal: Tolmanian maps.* We propose that the hippocampus contributes to planning by instantiating an episode-based map. During search, episodes are retrieved, modified, and combined to produce new plans as formally described in case-based planning (Borrajo et al., 2015). We term these maps “Tolmanian” because his latent learning paradigm must be solved using episodes rather than abstract representations.

Complicating matters further, other forms of planning that have received less attention in the literature do not appear to be hippocampally-mediated. For example,planning in chess depends on rules, perceptual patterns of board arrangements, and chunks of actions (Gobet, Retschitzki, & Voogt, 2004), planning a new a recipe depends on imagined physiological sensations related to taste, and planning a dance choreography depends on simulating novel motor combinations – none of which are normally associated with hippocampal function (Dayan & Berridge, 2014).

We suggest that it is unlikely that a single neural system represents all of the possible states and transitions associated with such a wide range of phenomena. Because long-term memories and cognitive maps both constitute sets of associations between items, we propose that planning problems are separated along the divisions of long-term memory: episodic, **semantic**, **Pavlovian** and **procedural**. As with the fable of the sages exploring different parts of the elephant (Fig. 2), each memory system represents a unique aspect of a more general map. In what follows, we survey this landscape by describing the unique mnemonic properties for each type of map and reviewing the evidence in support of them.

**Figure 2.**
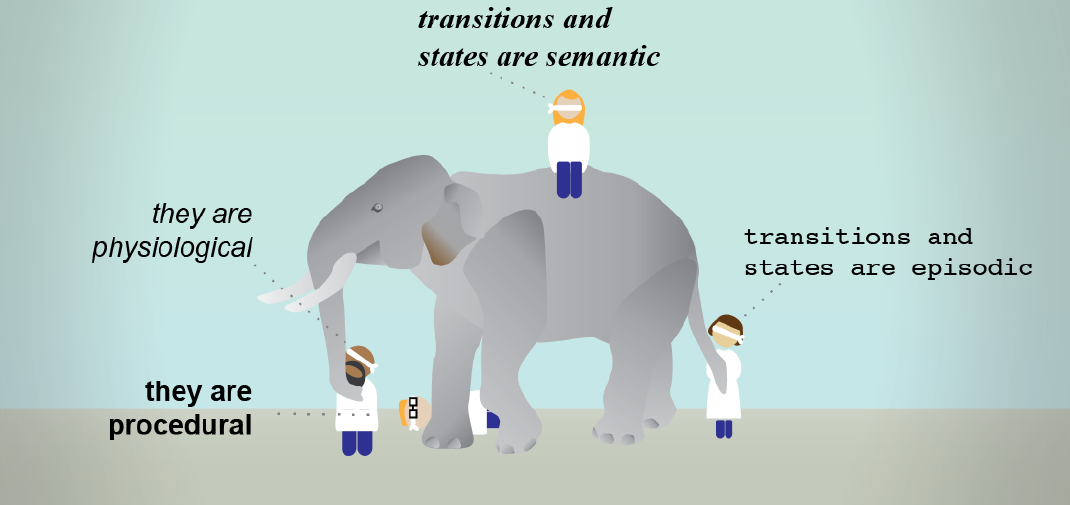
The Elephant and the Sages. In an old Indian parable, a group of sightless sages are asked to describe the appearance of an elephant based on touch. Because each person feels a different part of the animal, their descriptions are partial and idiosyncratic. Similarly, we propose that different long-term memory systems implement separate cognitive maps, each of which imposes unique constraints on how states and transitions are represented in planning problems: episodic (or Tolmanian) and semantic, procedural and Pavlovian (or non-Tolmanian) maps. As with the fable, integrating these descriptions into a unified representation presents a challenging computational problem. Original artwork by Gil Costa (Champalimaud Neuroscience Programme).

## A Framework for Hippocampal Function: Tolmanian Cognitive Maps

We propose that the hippocampus encodes an episode-based cognitive map, a hypothesis that reconciles its seemingly incompatible involvement in planning and episodic memory formation (Fig.1C). In our proposal, the hippocampal cognitive map consists of a web of episodes, or unique spatiotemporal trajectories. During search, episodes are retrieved, modified and recombined to produce novel action policies. We term models based on episodic memories as **Tolmanian cognitive maps** because Tolman’s *latent learning* paradigm emphasized aspects of episodic learning such as a short number of exposures to a maze (Tolman, 1932). By contrast, we term models based on other sources of memory as **non-Tolmanian cognitive maps**.

*What is the evidence for Tolmanian maps?* Evidence for hippocampal involvement in planning is extensive (Box 2). Crucially, this research has relied on paradigms that exercise elements of episodic function, reflecting Tolman’s seminal influence. Our framework also aligns with a nearly forgotten proposal of Ivane Berishtavili, who was a student of Pavlov (Tsagareli, 2015), that animals solve spatial latent learning problems based on *images* of the spatial layout – a concept that is reminiscent of episodes.

*How are states defined?* Consistent with the properties of episodic memory, Tolmanian states are rich in spatial detail and tagged to unique times and places: *yesterday, 5 p.m., sunny afternoon*. A sequence of states and transitions defines an episode.

*How are transitions defined* ? Transitions are unique spatiotemporal trajectories, short in temporal duration, that move between states deterministically – much like a *Vine* video (Hasselmo, 2012).

*How are the maps organized?* We suggest that hippocampal maps consist of connected clusters of related episodes, in line with existing research on spatial cognition (Han & Becker, 2014).

*Does the content reflect single or multiple experiences?* Each entry in the map refers to a single experience.

*How are they used?* Planning based on episodes can happen by retrieving and modifying single episodes, or by recombining multiple episodes. Case-based planning, a subfield of artificial intelligence, provides a formal description of how such a process could operate (Borrajo, RoubÍčková, & Serina, 2015; Hammond, 1986). In this framework, memories are retrieved and adapted to form new plans. For example, a plan to travel from campus to downtown could link episodes of “I took Bus 15 from campus to the Main Station” and “I took Bus 7 from the Main Station to downtown”, while ignoring goal-irrelevant details of the memories (such as, “I met Mr. Smith on Bus 15”).

*When are they used?* Episodic memory is advantageous for novel planning problems when there is little experience to form a general rule. The influence of episodes in planning decreases with increasing experience with the problem domain, enabling other long-term memory systems to take command.

## Non-Tolmanian Cognitive Maps: Semantic, Pavlovian and Procedural

In the next section we argue for the existence of cognitive maps that have received less attention in the literature: Semantic, Pavlovian, and Procedural maps. Although differing in specifics, these maps share in common preferentially non-spatial representational structures, and generalize information across individual instances.

### Semantic Cognitive Maps

Semantic memory comprises a variety of knowledge structures that abstract over instances (Ghosh & Gilboa, 2014; Tulving, 1985). Facts, such as “Ottawa is the capital of Canada”, which describe conceptual associations that are divorced from specifics and context, provide a quintessential example of a type of semantic memory. Another example is the statement “Napoleon lost at Waterloo”, which is a semantic concept except for the people who actually lived through the event. Other forms of semantic information include rules, scripts, categories, schemas, narratives, and statistical regularities (Ghosh & Gilboa, 2014). Because of its abstract property, semantic memory is a good candidate for encoding cognitive maps as they are usually understood (Gershman & Daw, 2017).

*What is the evidence for semantic maps?* Neuropsychological studies provide compelling evidence for semantic-based search (Irish et al., 2016; Irish, Addis, Hodges, & Piguet, 2012; Klein, Loftus, & Kihlstrom, 2002). Patients with semantic dementia, a neurodegenerative disorder affecting the anterior temporal and frontal cortices, are impaired at predicting abstract future events such as “What are likely medical advances of the next years?” (Klein et al., 2002). Yet their ability to retrieve past episodes is no different than that of control subjects. Furthermore, when asked to imagine “a one-off event in their future lives”, these patients tend to retrieve past episodes even when asked explicitly *not* to do so (Irish et al., 2012). In contrast, patients with hippocampal lesions can predict abstract future events (Klein et al., 2002); in a landmark study that revealed impairments in imagining details of a future day at a market, patients could still name appropriate market locations, even suggesting particular outdoor locations for sunny days (Hassabis, Kumaran, Vann, & Maguire, 2007). Recent rodent electrophysiology and human neuroimaging experiments suggest that the orbitofrontal cortex implements an independent cognitive map (Schuck, Cai, Wilson, & Niv, 2016; Wilson, Takahashi, Schoenbaum, & Niv, 2014), aligning with neuroimaging evidence that the ventral medial prefrontal cortex encodes schemas of semantic action structures (Ghosh & Gilboa, 2014). Finally, diary reports have identified an abstract mode of elaborating future scenarios (consistent with semantic memory), independently of a more visual and detail-oriented mode (consistent with episodic memory) (Stawarczyk, Cassol, & D’Argembeau, 2013).

*How are states defined?* Semantic states describe abstract concepts lacking in detail (Szpunar, Spreng, & Schacter, 2014). Because they do not necessarily depend on direct experience, semantic states can encompass larger domains of knowledge relative to episodic states, such as EUROPE.

*How are transitions defined*? Semantic transitions, like semantic states, are abstract, describing the consequences of general actions (e.g., “getting a pet”) rather than of specific actions (e.g., visiting a particular pet store). Semantic transitions can encompass different forms of knowledge. For example, statistical regularities, which represent the probability of transitioning between two states given an action, as derived from multiple exposures, constitute the fundamental mechanism underlying formal models of planning (Turk-Browne & Scholl, 2010). By contrast, rules that describe arbitrary, usually deterministic relations between states – such as “in chess, bishops move diagonally” – can be learned with a single exposure. Importantly, semantic transitions can be less constrained by temporal relations than episodic transitions. For example, the following transition has no obvious before-after: applying the action “What is the genus?” to the state “Chimpanzee” yields the state “Pan”.

*How are they organized?* Semantic maps should display preferential access to specific levels of abstraction, such that planning at the level of abstraction of the map should be faster and more accurate (Rosch, 1975). In other words, it would be easier to decide between getting either a dog or cat, rather than between specific dog or cat breeds, unless they are a dog or cat expert.

*Does the content reflect single or multiple experiences?* Each entry constitutes an abstract representation that is dissociated from individual episodes, even when learned in a single exposure. For example, “Tirana is the capital of Albania” can be learned with a single exposure to the fact without ever setting foot in the country.

*How are they used?* Because semantic representations naturally align with the concept of model states and transitions, they conform to the prescriptions of model-based reinforcement learning (Daw et al., 2005).

*When are they used?* Semantic cognitive maps are available for use when statistical regularities have been extracted from multiple individual experiences, or when rules apply to the decision problem. These predictions are powerful because they apply to a wide variety of states. For example, when parking a car every day in the same parking spot, semantic memories of the town layout are likely to inform how to return to the car from any arbitrary location. And when parking in an entirely new city, the instructions from a friend can inform how to reach it.

### Pavlovian Cognitive Maps

Pavlovian memory associates external states (such as a ringing bell) with physiological states (such as satiety), as well as between different physiological states (such as satiety and sleepiness) (Redish, 2013). It has been proposed that emotions constitute Pavlovian responses that manifest in spectrums of physiological and cognitive associations (LeDoux, 1995). Pavlovian maps would thus relate internal states to each other and to external variables. For example, Pavlovian maps could be used to predict the effects of food ingredients on gustation, enabling the creation of new tastes. Likewise, a model that mapped external situations to emotions could be applied by an actor to predict a feeling of sadness in an upcoming scene.

*What is the evidence?* Revaluation paradigms, such as devaluation due to pairing of an appetitive choice with nausea, first provided the evidence for planning (Balleine, Doherty, & O’Doherty, 2009; Daw et al., 2005) (Box 1). Further refinements of devaluation paradigms have indicated that these effects rely specifically on Pavlovian information. In particular, rats can rapidly modify their behavior on the basis of a previously learned association between a tone and the ingestion of a hypersaline solution. Although animals normally avoid the solution, under a novel condition of sodium depletion, they immediately chose it (Robinson & Berridge, 2013; Wirsig & Grill, 1982). Based on these experiments, it has been proposed that Pavlovian model-based reinforcement learning constitutes an independent form of planning, consistent with our proposal of independent Pavlovian maps (Dayan & Berridge, 2014; Pezzulo, Rigoli, & Friston, 2015).

*How are states defined?* Consistent with Pavlovian learning, Pavlovian states encode perceptual and physiological information (LeDoux, 1995). In contrast to other classes of cognitive map, these representations are non-declarative (semantic and episodic), poor in spatial detail (episodic), and not action-centric (procedural).

*How are transitions defined?* Transitions are of two types: from external states to physiological states (e.g., from a blinking light to satiation), and between physiological states (e.g., from satiety to sleepiness, hunger to anger, and so on). Although the transitions are not directly motor-related, they can bias action selection through Pavlovian-instrumental transfer (Talmi, Seymour, Dayan, & Dolan, 2008).

*How are Pavlovian maps organized?* This remains to be determined.

*Does the content reflect single or multiple experiences?* Both: Pavlovian representations can be learned from one or multiples exposures to an event (Keith-Lucas & Guttman, 1975; Rescorla & Wagner, 1972). For example, even a single exposure to a spoiled food item can associate the food with nausea (Welzl, D’Adamo, & Lipp, 2001).

*How are they used?* Pavlovian maps are used whenever planning involves physiological changes.

*When are they used?* Because Pavlovian maps can be acquired with one or multiple exposures, they can be utilized either immediately upon their initial acquisition or later in learning.

### Procedural cognitive maps

Procedural cognitive maps represent collections of related actions such as “going up to the third floor” and “getting to the airport” (Dolan & Dayan, 2013; Hamilton & Grafton, 2007). The maps concern intermediate levels of the motor hierarchy, between detailed representations of motor control like the positions and velocities of joint angles, and abstract semantic categories (Hamilton & Grafton, 2007). It has been proposed that planning can be carried out based on action sequences, on what is termed model-based hierarchical reinforcement learning (Botvinick & Weinstein, 2014; Dezfouli & Balleine, 2013). According to this formulation, action sequences are used to predict the end states of the sequences given the start states, without going through the effects of every single action.

*What is the evidence?* Despite provocative evidence from behavioral experiments (Charness, Reingold, Pomplun, & Stampe, 2001; Huys et al., 2015; Solway et al., 2014), the cognitive mechanisms underlying procedural planning are unknown. We predict that patients with damage to other memory systems would still be able to plan using procedural memory. For instance, as a thought experiment, we predict that a guitar player with impaired semantic, episodic and Pavlovian memory could still follow instructions to play a piece that started and ended in the hand position of a C minor chord.

*How are states defined?* We hypothesize that procedural states are motor-centric, integrating the state of the motor system with the sensory consequences of actions (Crump, Logan, & Kimbrough, 2012), akin to what have been termed elsewhere as *ideomotor representations* (Hommel, Müsseler, Aschersleben, & Prinz, 2011). An example would be “holding the guitar before playing”. Procedural states have the capacity for varying degrees of abstraction (Botvinick & Weinstein, 2014). Furthermore, they are not declarative, even when highly complex.

*How are transitions defined?* Procedural transitions map the beginnings to the ends of a sequences of actions without encoding the intervening steps (Botvinick & Weinstein, 2014).

*How is the map organized?* Procedural maps are organized according to a strict hierarchy of different levels of abstraction (Logan & Crump, 2011). Thus, the representation of “going to the airport” could be composed of action sequences related to “going through McKenzie Road”, “taking highway 17” and “taking Airport Road”, allowing the planning mechanism to access “going to the airport” without searching through the three sub-sequences (Botvinick & Weinstein, 2014).

*Does the content reflect single or multiple experiences?* Procedural maps encode information accumulated across multiple experiences, in accordance with the slow acquisition of habits (Daw et al., 2005; Dolan & Dayan, 2013).

*How are they used?* Procedural maps allow for the compositional sequencing of actions to achieve a higher-level plan, like combining short musical sequences in order to compose a longer score.

*When are they used?* Procedural maps take longer to acquire than episodic and semantic maps (Botvinick & Weinstein, 2014). However, once learned, their strict hierarchical organization – where upper levels are blind to transitions between lower levels – allows for efficient planning (Botvinick & Weinstein, 2014; Solway et al., 2014).

## Concluding Remarks: Reconstructing the Elephant

We have proposed that planning relies on parallel cognitive maps, each of which is mediated by a specific form of long-term memory associated with a distinct neural system: episodic, semantic, Pavlovian and procedural. Each of these maps represents the decision problem according to the language it speaks, with varying degrees of abstractedness, and different dimensions. This framework has three important consequences: it alleviates the burden of a single map representing the myriad of possible states that occur in most decision problems, it reconciles apparently opposite views of hippocampal function by placing the hippocampus as the source of episodic planning, and it highlights an essential, and often overlooked, role of other long-term memory systems in planning. Yet, the proposal raises as many questions as it addresses.

In particular, this framework raises the question of how the planning systems communicate with one another (Fig 2). Are their outputs integrated in a single brain area? The results of neurological and animal lesions studies are telling in this regard. Such a brain area could not be neocortical, because rats with neocortical ablations display physiological latent learning (Wirsig & Grill, 1982), thus eliminating from the discussion neocortical areas associated with planning: medial and lateral orbitofrontal cortices (McDannald et al., 2012; Wikenheiser & Schoenbaum, 2016), dorsolateral prefrontal cortex (Smittenaar, FitzGerald, Romei, Wright, & Dolan, 2013), and premotor areas (Kornysheva & Diedrichsen, 2014). Outside of the neocortex, hippocampal lesions do not impair planning based on semantic memories, ruling out this system (Irish et al., 2016; Klein et al., 2002). This process of elimination implicates subcortical nuclei, especially brain areas previously associated with planning like the amygdala and striatum (Daw & Dayan, 2014; Deserno et al., 2015; Prévost, McNamee, Jessup, Bossaerts, & O’Doherty, 2013), but this possibility remains to be investigated.

Alternatively, the plans could be integrated closer to the action production stage, with different plans competing for control over the motor system (e.g., (Holroyd & Coles, 2002)) similar to discussions of controlled vs. automated and dual-system theories of behavior (O’Reilly, Noelle, Braver, & Cohen, 2002). For example, whereas a semantic model might promote behaviors that avoid caloric foods, a Pavlovian model might do the opposite, giving rise to seemingly irrational actions as different systems vie for control. These plans must also be integrated with habitual behaviors: Whereas previous theoretical treatments have characterized decision making as a competition between two systems – one model-based (for plans) and one-model free (for habits) (Daw et al., 2005) (Box 1) – our framework potentially increases the complexity of this problem by a factor of four, with separate pairs of systems for each category of memory. To complicate matters further, procedural and Pavlovian model-free systems may be easy to imagine, but semantic and episodic model-free systems seem less so.

Clearly, researchers have only just started to chart the world of cognitive maps. Although much remains to be learned, we hope that our efforts have defined its largest continents.

## Glossary

**Abstraction:** Computational term indicating the degree to which a representation encompasses other subordinate representations. Abstract representations apply to multiple items, such as the semantic concept “take a means of transportation”, which encompasses “take bus”, “take bike”, and others. Episodic representations lie at the other extreme, such as “take bus 15 on Thursday evening”.

**Cognitive map:** Cognitive maps have historically referred to an internal construct that allows for spatial latent learning. More recently it has been equated with the concept of a model. Here we generalize the concept to any representation of the consequences of actions that allows for planning.

**Episodic memory:** A form of declarative memory for experiences that preserves information about unique times and places.

**Hippocampus:** An allocortical structure within the medial temporal lobe that supports the formation, consolidation and retrieval of episodic memories.

**Model**: A computational representation of consequences of actions, composed of a one-step transition function that specifies the probability of transitioning between states given an action, and a reward function that predicts the amount of reward accrued following a transition.

**Model-based reinforcement learning**: The set of computational processes involved in learning and utilizing models of decision problems for planning.

**Non-Tolmanian cognitive maps**: Semantic, Pavlovian and procedural cognitive maps.

**Pavlovian memory**: The set of learned associations between external states and somatic or physiological states, including emotional memories, such as an association between a ringing bell and satiation.

**Planning:** In model-based reinforcement learning, the process of searching through the consequences of action sequences before the actions are taken. It is equivalent to the concept of goal-directed behavior.

**Preplay/replay**: Preplay refers to the sequential activation of place cells corresponding to potential routes in a maze. Replay refers to sequential activation of place-cells corresponding to past experiences.

**Procedural memory**: Following precedent, we equate procedural memory with the model-free system and habits (Dolan & Dayan, 2013), although it is likely to encompass other types of memory as well (e.g., cerebellar).

**Semantic memory:** A form of declarative memory for abstract knowledge of the world, irrespective of the context in which it was learned (e.g., facts, rules and statistical regularities).

**Search**: The process of sequentially probing the transition and reward functions in a model, which is the formal equivalent of mentally simulating the consequences of actions. Prospection, future-thinking, simulation, preplay, vicarious trial-and-error and foresight can be conceived as search.

**Tolmanian cognitive map:** Episodic or instance-based cognitive map consisting of a set of episodes, each of which corresponds to a unique spatiotemporal trajectory. We hypothesize that this type of map underlies planning in Tolman’s latent learning tasks.

**Transition**: Computational term denoting the change from one state to another state after performing an action.

## Box 1: Planning

Planning, or goal-directed behavior, is a mode of decision making in which the values of actions are determined by searching through potential courses of action in order to evaluate their consequences before the actions are taken (Geffner & Bonet, 2013; Redish, 2013). The search process is based on a model of the decision problem that represents the consequences of actions. The subject of model-based reinforcement learning (Daw et al., 2005) describes both the process of learning the model, which is formally defined by a matrix of abstract state-action transitions, and using the model in order to plan. In contrast, no search takes place in Pavlovian and habitual decision making (Daw et al., 2005; Dolan & Dayan, 2013), the latter of which is sometimes referred to as model-free reinforcement learning. A hallmark of planning is rapidly modifiable, flexible behavior, in contrast to habitual behavior, which can be changed only slowly.

Planning occurs in a variety of domains: social settings, where the model predicts the behaviors of other people (Erev & Roth, 1999), intertemporal choice, where the model predicts the future effects of saved rewards (Kurth-Nelson, Bickel, & Redish, 2012), problem-solving, where the model predicts abstract outcomes according to a set of rules (Newell & Simon, 1972), spatial navigation, where the model is a spatial map (Spiers & Maguire, 2008), and even physiological activity, where the model predicts upcoming physiological or emotional states (Dayan & Berridge, 2014).

Several empirical paradigms have been developed to reveal evidence of planning. The classic approach depends on showing that a subject can produce novel, adaptive behaviors without it having enacted exactly the same behavior previously (Balleine et al., 2009; Tolman, 1948). For example, in spatial latent learning paradigms, a rat is initially allowed to explore a maze in the absence of any reward; subsequently, when the rat is presented with a food reward at a specific maze location, it immediately returns to that location on the following trial, revealing latent knowledge of the maze structure that it uses for navigation (Tolman, 1948). And in devaluation paradigms, evidence of planning is seen in an immediate change in behavior that follows a sudden change to a choice outcome. For instance, rats immediately eschew choices that lead to poisonous outcomes, even when these same choices were rewarding in the recent past (Balleine et al., 2009). Finally, a major innovation in the study of planning was introduced with the human “two-step” task, which repeatedly tests for knowledge of transitions and rewards during a single experiment (Daw, Gershman, Seymour, Dayan, & Dolan, 2011). The paradigm reveals rapid changes to choice probability based on recent updates to expected reward value.

## Box 2: Hippocampus, Episodic Memory and Planning

## Episodic Memory

A role for the hippocampus in the formation and retrieval of episodic memories is well-established (Dickerson & Eichenbaum, 2010; Moscovitch, Cabeza, Winocur, & Nadel, 2016). Episodic memory is a form of declarative memory that encodes information specific to the time and place of acquisition and that is usually rich in perceptual detail (Tulving, 2002). Damage to the hippocampus impairs the acquisition, retention and retrieval of new episodic memories while sparing semantic memories and the gist of previous events (St-Laurent, Moscovitch, Jadd, & McAndrews, 2014; Viard, Desgranges, Eustache, & Piolino, 2012). Functional neuroimaging corroborates the neuropsychological findings. For example, hippocampal activation is positively correlated with the amount of detail of recalled events (Cabeza & St Jacques, 2007). The episodic network also overlaps with spatial tasks (Spreng, Mar, & Kim, 2009), with greater anterior hippocampal engagement for memories of events that occurred on a larger scale (Poppenk, Evensmoen, Moscovitch, & Nadel, 2013).

## Planning

A second body of research links hippocampal function with planning by positing that the hippocampus encodes a cognitive map (Wikenheiser & Redish, 2015). This proposal is based on the discovery by O’Keefe and Dostrovsky of *place cells*, which are hippocampal neurons that respond selectively to the current position of the animal (O’Keefe & Dostrovsky, 1971). More recently, evidence that place-cells serially code for positions ahead of the animal at decision points in a maze, termed *preplay,* appeared to close a circle between planning, the hippocampus and the cognitive map (Wikenheiser & Redish, 2015).

In line with the findings in rodent studies, hippocampal damage in humans impairs imagining new experiences (Addis & Schacter, 2011). For example, patients with hippocampal damage provide impoverished descriptions of a future day at a market, neglecting to mention the specific merchants, shops, and sounds of a marketplace (Hassabis et al., 2007). Hippocampal damage and the inability to imagine future experiences is also positively correlated with impairments in decisions involving future outcomes (Palombo, Keane, & Verfaellie, 2015). Human functional magnetic resonance imaging studies have corroborated these findings; for instance, instructions to imagine future scenarios – such as, “imagine a day at the market” – activate the hippocampus (Addis & Schacter, 2011). Recent work under the framework of model-based reinforcement learning has further highlighted a role for the hippocampus in planning (Balaguer, Spiers, Hassabis, & Summerfield, 2016; Bornstein & Daw, 2013; Daw & Dayan, 2014; Doll, Duncan, Simon, Shohamy, & Daw, 2015).

